# Enterotype-like microbiome stratification as emergent structure in complex adaptive systems: A mathematical model

**DOI:** 10.1101/402701

**Authors:** Miguel Ángel Martín

## Abstract

**Background:** Metagenomics has provided valuable insight into human gut microbiome composition together with its structure and function. Studies suggest that the adult gut microbial community possess a certain amount of stability and resilience. The architecture of host-microbial symbiotic states suggested by microbiome clustering findings and co-occurrence functional networks might be involved in these properties. Models for understanding the underlying structure (including enterotypes) and derived properties are in demand by researchers.

**Principal findings:** We propose a simple random function system as a model approach of the adaption and self-organization of the microbiome space when fostering the optimal functioning of the system. The construction of this model is based on key facts of microbiota functioning reported in recent studies. We aim to demonstrate the existence of a probability distribution as a microbiome attractor resulting from an intermittent adaption process. Its mathematical structural properties explain the stability of gut microbiota and its ability for restoration after occasional perturbations. The model is consistent with microbiome clustering results and provides precise mathematical meaning to the gradient among enterotypes previously reported. The model also explains how intermittent perturbations, such as long-term dietary patterns, might affect the microbiome structure, and these results are consistent with previously reported experimental results.

**Conclusions/significance:** These mathematical facts implied by the model unveil an underlying mechanism that may explain gut microbiome structure and related experimental findings. Within this framework, stability and resilience properties of human gut microbiota are explained as a consequence of the model.

## Introduction

The human gut houses billions of microbes and composes the most densely populated microbial ecosystem in the human body, encompassing greater than 1000 species. Microbial community composition changes rapidly in early childhood. The microbial community is fully developed by the age of approximately 3 years, and healthy adults appear to have a unique and relatively stable gut microbiota. By analyzing the 16S ribosomal RNA-encoding gene, metagenomics studies have reported a great deal of information on phylogenetic composition, bacterial functions and ecological dynamics in gut microbiota. Several studies support the hypothesis that although gut microbiota varies widely among individuals at the phylogenetic level, metabolic genes and pathways are conserved, accounting for a “core microbiome” [1]. In 2011, a clustering analysis of metagenomics fecal samples of individuals from six different nationalities [2] led to the identification of three robust clusters or “enterotypes” that do not seem to correlate with nationality, gender or age. Each enterotype was characterized by variation in the levels of three respective genera (“drivers”) whose abundances are negatively correlated. Enterotype 1 is dominated by the genus *Bacteroides,* enterotype 2 is dominated by *Prevotella* and enterotype 3 by *Ruminococcus*. Additionally, the drivers are the hubs of three concurrent ecological networks, indicating different combinations of microbial trophic chains that generate energy from fermentable substrates are available in the colon [2].

Later studies recovered the three enterotypes [3-6], whereas others found that the human gut microbiome could be separated into a different number of optimal clusters [7], which has generated an intense debate among the scientific community. Different authors noticed the weak separation between clusters and proposed a gradient-based interpretation, suggesting the substitution of discrete clusters by “enterogradients” [8-12]. A review together with a detailed discussion on enterotype and gut stratification issues is presented in [13].

Underlying the enterotype debate, there is an implicit recognition of organization structure in the gut microbiome that is also suggested by the relative stability of adult microbiota and *resilience*, which is the tendency to return to a previous state after periods of disturbance. Microbial communities may be considered complex adaptive systems. From interactions between microbes and with the environment, the global community may exhibit emergent properties that are not present in individual microbes [14]. Self-organization, which is the tendency of a large dissipative system to form optimal functioning states [15,16], has been invoked as an explanation of biological complexity [17]. Gradient adaptation is a key component of the formation of optimal states [18]. However, gradient adaption principles that apply to a single trophic compartment are no longer generally true for “complex ecosystems,” i.e., when greater than one trophic level is present [19]. The basic mechanisms underlying adaptation and self-organization remain unclear, and different approaches, including mathematical methods, need to be better integrated [20]. In addition, different models are lacking as reference to describe alternative interpretations of the enterotype concept and the assumptions that underlie these different interpretations [12]. This concept is related to both demands, demonstrating both how mathematics provide insight for understanding the “roots” of the complexity derived from certain adaptation mechanisms and how a mathematically based model may help to better understand the structure and properties of the gut microbiome.

## Methods

### The model

A simple adaptive dynamical model is formulated to account for underlying mechanisms leading to gut microbiome clustering structure and stability and resilience properties. Based on basic biological facts reported in [1,2] and related studies mentioned above, the model is built with a focus on the adaption of gut microbiome to a set of metabolic genes thought to promote key metabolic functions without delving into the enterotype debate about the number of driver taxa and co-occurrence networks or considering the different combinations of microbial trophic chains involved.

### Adaption considerations

Basic ingredients of adaption theories developed in [16,18,19,20] were used. Optimal functioning in a network structure should correspond with “optimal” metabolic gene catalogues. Mutations and genetic rearrangements enable an organism to adapt to environmental conditions. An adaptive process is should also operate in the space of gene sequences by means of a gradient adaption towards metabolic genes. An intermittent random action would catalyze certain gene configurations. The system should meet recurrent demands in more than a single direction, which is a main component of the core of our model. A loose mix of randomness and determinism coexistence (as a type of “chance and necessity” [21]) should lead to optimal microbiome states. In the process, a variety of gene pathways indicating “flow gradients” towards primary metabolic gene configurations are generally expected.

### Assumptions of the model

Let us denote the microbiome code space by *X*. Although changing distance metrics or clustering algorithm yields a different number of clusters even for the same data set [7,13], an abstract conceptualization of gene sequences as points in a mathematical metric space might be useful in modeling the gut microbiome. Several distances (Jensen-Shannon, Unifrac-weighted, Bray-Curtis, and Euclidean) [22] permit viewing of (*X*, *d*) as a mathematical metric space. To fix ideas and for visualization purposes, we shall propose that *X* is the bidimensional Euclidean space *IR*^2^ provided by the Euclidean distance. Due to the multidisciplinary nature of this work, we shall use a sufficiently precise presentation albeit avoiding strict mathematical formalism, which can be observed in mathematical references.

We assume the following:

A1: A family of random functions {*φ*_*ϑ*_: *ϑ* ∈Ω} acting on the state space *X* adapting gene configurations to those of metabolic genes exists.

A2: These functions act in an intermittent manner as they are driven randomly by the request of optimal functioning of the system. A probability distribution *μ* on Ω controls the intermittent regime required to optimize system function. This distribution is supposedly determined by the host and environment conditions.

### From gene pathways to gene abundance probability distribution

The idea is that a Markov chain drives the “gene dance” as follows. If the chain starts at *x*_0_ ∈ *X*, it moves by choosing at random *ϑ* from *μ* and going to 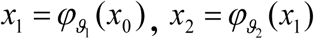. and inductively 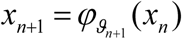, where *ϑ*_1_, *ϑ*_2_,…are independent draws from *μ*. Thus, the successive configurations do not depend of the past.

Some might pose a question about the features of gene pathways and whether the sequences of configurations obtained from initial configurations in the space X draw, in some sense, a long-term configuration structure of the gene “abundant distribution (measure)”.

Theory provides an affirmative answer.

Theorem 1: Under the hypothesis above, if the functions {*φ*_*ϑ*_: *ϑ* ∈Ω} are contractive (“by average”), then there is a unique stationary probability distribution *π* on *X* with
*P(x*_*n*_ *∈A)→π(A)* as *n→∞*

which does not depend on the starting point *x*_0_ ∈ *X*. Here, *π* (*A*) indicates the gen abundance in any subset *A* of the gene configuration space.

The proof of this theorem is obtained from Diaconis and Freeman [23]. The expression “by average” is stated in a technically precise manner.

The relevance to our model is derived from the fact that the adaptation is expected to be done by reducing the distance of gene configurations to those of metabolic genes. Thus, in mathematical terms, the functions in A1 should be contractive (statistically). As a consequence, under the frame of this model, the gut microbiome appears as a well-defined structure whenever the random function system and the probability distribution *μ* are given by the host and the environmental conditions.

As an example, the reader may think that the chain is defined by affine random functions in the 2-D Euclidean space:

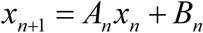

where *A*_*n*_ are identically distributed in a 2×2 random contractive matrix, and *B*_*n*_ are 2×1 vectors randomly distributed on gene configuration drivers.

Theorem 2. Under certain general conditions, for any starting point, the convergence of *P(x*_*n*_ *∈A)→π(A)* occurs at an exponential rate. This notion is more precisely stated below.

For any *n* the law of *P*_*n*_ (*x*,.) is given that *x*_0_ = *x* and ρ is the Prokhorov metric used as distance between two probability distributions. Then, *ρ*[*P*_*n*_ (*x*,.),*π*]≤ *A*_*x*_ *r* ^*n*^, and *A*_*x*_ and *r* serve as constants (0 < *r* < 1). These bounds apply to all *n* values and all starting points *x* (see [23]).

Within the frame of this model, this theorem actually explains the resilience property of gut microbiota, namely, to tendency to return to the previous stationary state if change results from some disturbance. Under the model hypothesis, if external forces occasionally modify the probability distribution, restoration occurs at an exponential rate.

### A simplified working model

Let *φ*_*i*_ be contractive functions *i* = 1,2,…*N*, i.e., *dist*(*φ*_*i*_ (*x*),*φ*_*i*_ (*y*)) < *dist*(*x, y*). Given that the possible utility of the model does not rely on the value *N* or the remaining values, we shall suppose *N* = 3. Given that *φ*_*i*_ are contractive mappings, they have invariant or fixed points that represent the respective gene configuration drivers. To provide a more graphic illustration, the reader may assume that *φ*_*i*_ represents fixed functions instead of random functions and that they are affine maps.

*φ*_*i*_ (*x*) = *a*_*i*_ + *b*_*i*_ Here, each *a*_*i*_ is a 2×2 contract, whereas *b*_*i*_ is a 2×1 vector.

Let *p*_*i*_ probabilities or weights (*p*_1_ + *p*_2_ + *p*_3_ = 1). These ingredients specify a Markov chain as follows: starting at *x* with the probability the chain proceeds by choosing at random *i* with probability *p*_*i*_ and moving to *φ*_*i*_ (*x*). The set {*φ*_1_, *φ*_2_, *φ*_3_; *p*_1_, *p*_2_, *p*_3_} is called an *iterated function system* (IFS). Here, *φ*_*i*_ represents the fitness functions and does not necessarily have a globally applicable value but might have individual-specific host-like character. On the other hand, the probabilities *p*_*i*_ reflect the frequency of the demands that the metabolic functioning requires. In a simulation with usual electrical network functioning systems, the probabilities *p*_*i*_ might represent automatic control, which allows regulation of adequate flow of energy through the network at any given point in the system. However, this discussion point falls beyond the purpose of this abstract model.

Next, an operator *X* is defined as a type of “adaption operator” acting on the microbiome phase state *X* and is defined as follows:

*H* (*X*) = *φ*_1_ (*X*) ⋃*φ*_2_ (*x*) ⋃*φ*_3_ (*X*), which transform the space *X* in the union of three respective transformed phase states and *H* ^*n*^ (*X*) = *H*(*H* ^*n*-1^(*X*)) the n-iterate.

Theorem 3: *H* ^*n*^ (*X*) converges (in the “visual” metric, technically described by the Hausdorff metric) to a configuration attractor *A*, which verifies the self-similarity property:

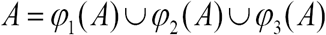

The proof of this result is provided by Hutchinson [24].

Moreover, the attractor *A* is the support of the probability distribution *π* whose existence is guaranteed by theorem 1. For simulation purposes, we’ll use the following Eltońs theorem:

Theorem 4. Under the same hypothesis of theorem 3, for any subset *B*, if δ _*x*_ (*B*) = 1, δ _*x*_ (*B*) = 0 if *x ∉B,* then 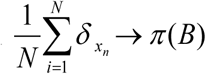.

See [25].

This approach might be relaxed in several directions to yield a better fitting model regarding the gut microbiome scenario. In particular, the functions may be contractive random functions instead of fixed function. The probabilities *p*_*i*_ may be random numbers, and the condition Σ *p*_*i*_ = 1 may be replaced by the average value of the sum *E*(Σ *p*_*i*_) = 1. Additionally, the adaption operator may be iteratively transformed to any starting random probability distribution, leading to a random probability distribution as a model of gene abundance distribution (see [26] for mathematical details).

### Simulation

Theorem 4 allows the following simulations.

If 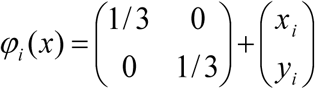, then 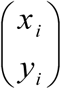 and *i* = 1,2,3 are the respective coordinates of an equilateral triangle.

The attractor is the so-called Sierpinsky attractor (Fig 1), which we shall use as schematic example in the Discussion section.

**Fig 1.**
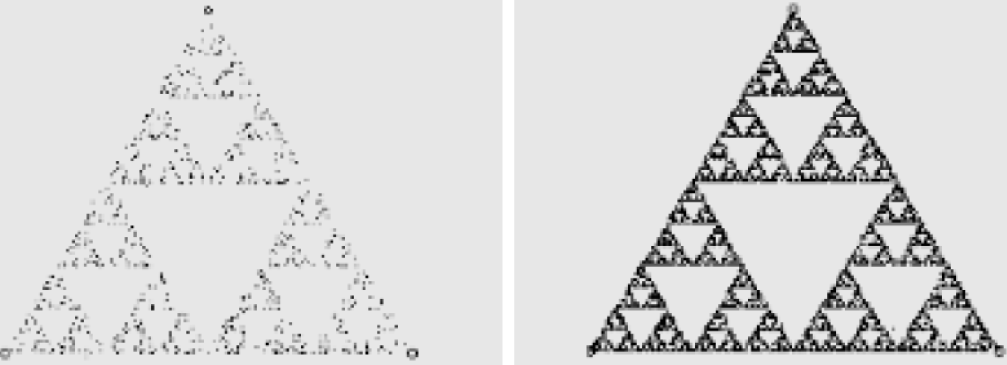
Simulation of the Sierpinsky Attractor with Equal *p*_*i*_’s 10.000 Iterations.

Using different systems of affine functions, one may obtain much more sophisticated attractors (see Fig 2).

**Fig 2.**
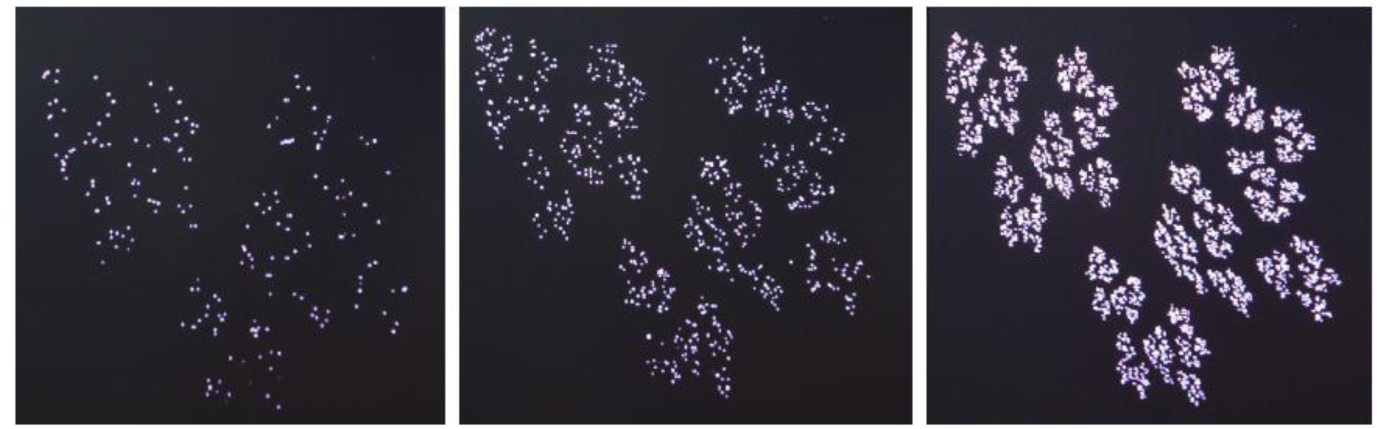
IFS Attractor by Implementing an Increasing Number of Iterations.

## Discussion

### Microbiome stratification under the modeling

According to this model, the asymptotic fractal probability distribution is supported on the “self similar” attractor *A*=*φ*_1_(*A*)∪*φ*_2_(*A*)∪*φ*_3_(*A*), which may be considered as the “core microbiome”, and their components, which may be considered as the basic “skeleton” formed by the respective basins of attraction around the driver taxa. Such components would be considered theoretical models for the clustering structures found in enterotype literature [2]. The recurrent application of the above self-similar equation for consecutive *n* = 1,2,… would lead to an hierarchical fractal type structure and abundance distribution measure supported in the rescaled copies **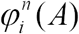,** which would represent the flow gradient in the direction to each driver taxa *j* = 1,2,3. This notion is consistent with authors proposing the “enterogradient” paradigm instead of discrete enterotypes [12]. The enterogradient draws a hierarchical stratified structure using basins of attraction or “halos” surrounding the drivers. The different levels in the network structure should correspond with different levels in metabolic genes. At any level, the model reveals substructures resembling the entire structure of the attractor *A*=*φ*_1_(*A*)∪*φ*_2_(*A*)∪*φ*_3_(*A*). This aspect would agree with gut stratification features reported (see [13] and references there in).

An interesting issue arises regarding how gene pathways evolve under the model hypothesis. A mix of chance and determinism guides the genes’ “dance” under this restricted modeling, producing a chaotic behavior along the entire journey. The points *x*_*n*_ follow a chaotic behavior in such a manner that they jump consecutively at random through the components 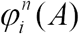 but surprisingly define a precise mass distribution (in probability distribution terms) with independence of the starting point *x*0. This is a key type of behavior derived from the existence of random intermittencies that push gene configurations towards more than one direction. In addition, repeated iterations in one single driver direction, which act to separate each gene, would explain the existence of discrete clusters. Thus, the random intermittent alternation leads to different hierarchically stratified structures surrounding the driver configurations.

Fig 3 illustrates the action of the adaption operator *H* ^*n*^ (*X*), demonstrating how the space X is iteratively reduced. Thus, microbiome configurations are confined to the attractor A as a type of “global flow” towards the drivers.

**Fig 3.**
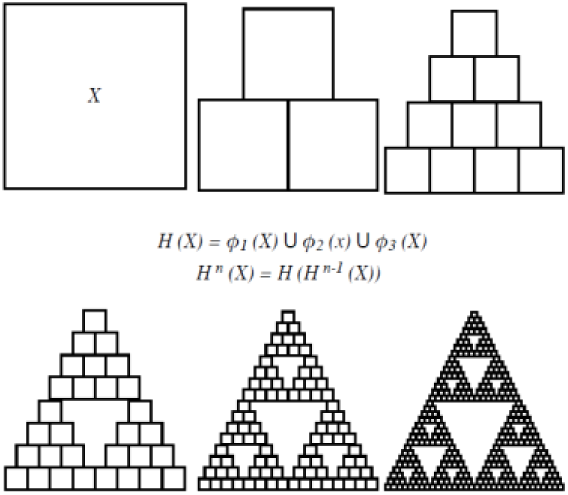
Schematic Illustration of the Action of the Adaption Operator.

Fig 4 suggests the possible compatibility of the microbiome stratification with co-occurrence of the hierarchical structure of respective metabolic networks. However, this point is mere speculation.

**Fig 4.**
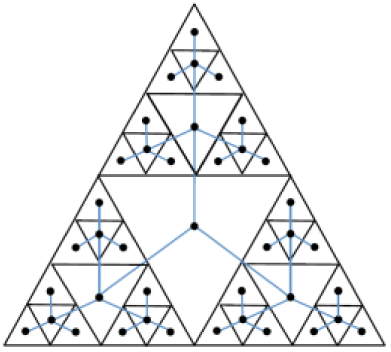
Schematic Illustration of Co-Occurrence Networks.

Interestingly, regarding basins of attraction [A=φ1(A)∪φ2(A)∪φ3(A)], the 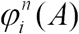 may overlap, which also is consistent with different studies where weak or fuzzy separation of clusters were reported ([13] and references there in).

When a given *p*_*i*_ is fixed, one of the two remaining probabilities is greater, whereas the other probability is lower, which should lead to the negative correlation of abundances in the enterotype clusters reported [2].

### Microbiome attractor as optimal entropy state

The complexity issue is another interesting point related with the proposed model. Indeed, the model proposed would explain the recognized stability of healthy human gut microbiota. The fractal attractor A as a model of probability that measures gut microbiome configuration is a type of scale-free structure that commonly appears in self-organized structures. Optimal principles as maximum entropy principles are very popular for characterizing optimal stable states in ecosystem functioning [27,28]. Given that entropy has different meanings (thermodynamic, information, or statistical) leading to different functional forms [29], our model is concerned with the Shannon Information Entropy, which may be used to characterize the attractor. If the corresponding abundances are *μ*(*φ*_*i*_ (*A*)) = *p*_*i*_ for *i* = 1,2,3, the Shannon Information Entropy [29] defined by 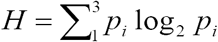 provides *p*_*i*_ log_2_ *p*_*i*_ = 0 if *p*_*i*_ = 0.

Fractal type structures produced by iteration schemes of the type used in our adaptive modeling, can also be inferred via the maximum entropy principle, which are systems characterized with a constant average generating information [30]. In fact, in our case, the Information Entropy of the probability distribution measure is computed at the scale level 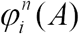, which constantly exhibits *self-similarity entropy*. The stability of natural fractal structures would then be ensured by the variation principle from which they were derived, and fractality would be a trend resulting from an economical method of achieving stability [30]. According to this observation, it is interesting to note that it would mean that discrete enterotypes would not guarantee the stability that healthy gut microbiota exhibit; stability requires a certain level of complexity (see [20] and references there in). Another reason for this intrinsic robustness is that the probability distribution model thrives on randomness. A certain amount of random perturbations are included in the growing process, facilitating the formation of the stable structure rather than hindering it. Finally, as indicated in section 2.3, Theorem 2 supports the resilience property in a mathematically based manner. Although this property has been used as a “stylized fact” or invoked on the basis of unclear arguments, here appears as a theoretical result within the frame of the model.

Information Entropy has been used to characterize stable ecosystems for a long period of time given that optimal ecosystem function is characterized by a rather well defined entropy value [31]. This finding is consistent with the existence of a limited number of well-balanced host-microbial symbiotic states in gut microbiota [2].

The model would also explain why long-term interventions, such as long-term dietary patterns, might influence in the attractor features given that such patterns might influence the activation of different trophic networks, which is reflected by the *p*_*i*_ values in the model.

### Prospective future research

Remarkably, given a target cluster image formed by a collection of points, theoretical results on IFS theory allows the possibility of identifying a set of contractive functions such that the collection of points {*x*_1_, *x*_2_ *x*_*n*_} forms a reasonable likeness of the target with high probability. This technique is widely used for image compression [32,33]. Such a possibility might potentially have a great number of applications in the modeling and simulation of gut microbiota function.

A second issue addresses the quantitative characterization of gut microbiota from the knowledge of relative abundances of driver taxa and the correlation with different functional properties. This is a common problem in hierarchically arranged multicomponent systems, i.e., soil, where the distribution of millions of particle sizes definitely influences its functional properties and the life developed within it. Although the soil scenario might seem different from the gut microbiota system, both systems share surprising coincidences with respect to taxonomy jobs. Interestingly, the characterization of soil texture has traditionally been made by reporting the mass percentages of only three size fractions (clay, silt and sand) despite being formed by such a complex mix of millions of particle sizes [34]. Depending on these percentages, soils have been classified in “textural classes” in a surprisingly similar manner given that a number of different ecotypes have been reported in microbiome studies [35]. Functional properties, such as soil water content or bulk density, exhibit a surprisingly excellent correlation with the IE simply computed using three fraction contents [36,37]. One possible reason for such a correlation would be that those functions depend on packing arrangement of particles. In gut microbiota, the correlation is derived by the fact that H might indicate a type of “packing network arrangement”. One may wonder whether a similar job might be performed in microbiota, which would require a significant amount of data collection similar to that noted for soil.

Finally, the simulating power of mathematical models constructed with these and new ingredients might represent a promising avenue to explore.

## Conclusions

The adaption of microbiome configuration to promote primary metabolic functional networks is modeled by means of a random function system that operates in an intermittent (random) manner on the microbiome configuration space. The gradient adaption to optimal functional states suggests that the functions should have a contractive action “by average”. The model demonstrates how the intermittent random action generates well-defined hierarchical complex probability distributions in the microbiome configuration space. The model helps to interpret the term “enterogradient” used in the literature as opposed to “discrete enterotypes”. The model also helps to understand gut microbiota as a complex system, thus providing a theoretical explanation of its resilience and stability and the possible influence of long-term dietary patterns in microbiome configuration.

The model suggests future research in several directions, including the identification of particular function systems to model target clusters. In addition, this information could be used for simulation purposes or as entropy parameters in microbiome characterization and correlations with functional properties.

## Acknowledgements

The help received from C. García-Gutierrez, I. Atienza and Alberto GonzÁlez in different aspects concerning the making of figures, is kindly acknowledged.

